# Rickettsia β-peptide Reactivity with Immune IgM

**DOI:** 10.1101/2024.06.11.598527

**Authors:** Lee Fuller, Susan Ordanza, Julia Reyna

## Abstract

Spotted Fever Group (SFG) Rickettsia species maintain an s-layer with two major outer membrane proteins (OmpA and OmpB) aligned within a lipopolysaccharide (LPS) matrix external to the bacterial outer membrane. While the two typhus group species (R. typhi and R. prowazekii) maintain a similar s-layer containing only OmpB with its own specific form of LPS. A major component of these two types of OmpB is a specific and highly conserved beta peptide (β-peptide) non-covalently attached to the respective OmpB passenger domain. This β-peptide is initially translated as the c-terminus of the 168 kDa polypeptide and initially forms a membrane pore to allow the larger passenger domain to exit the bacterial cytoplasm through the outer membrane. Prior to the full exit of this passenger domain from the cytoplasm the long polypeptide chain is cleaved by a peptidase, leaving the c-terminus remaining as a beta-folded membrane pore and the N-terminal passenger domain exiting to the space between the bacterial membrane and the s-layer. The fate of this pore structure and the attachment of β-peptide to the OmpB continues to be examined and will be discussed as another protective component of the Rickettsia. An EIA IgM assay has been developed to utilize this antigen for clinical diagnostic use to accurately detect acute rickettsial infection in testing labs and lead to the availability of rapid test (lateral flow) formats for more immediate testing.

## Introduction

The vast majority of Rickettsia serology testing worldwide is still performed using IFA slides where 1) the results strongly reflect reactivity to SFG and TG (typhus group) lipopolysaccharides (LPS), 2) the Rickettsia species is assumed to be the most prevalent one locally and 3) the IgM reactivity is too often false-positive due to these LPS antigens. This LPS reactivity is non-specific in that it is like that produced by the neonate against the maternal microbiome and is maintained for life. This reactivity is termed “Rickettsia reactive” instead of “Rickettsia specific”, which is specifically induced by a Rickettsia infection.

Rickettsia OmpB (sca5) belongs to a family of proteins called autotransporters, which have a modular structure including an N-terminal secretion signal, central passenger domain, and C-terminal β-peptide (βp). OmpB is initially translated as a 168-kDa polypeptide and is cleaved to yield a 120-kDa surface-exposed passenger domain that remains loosely associated with the outer membrane and a 32-kDa integral βp. The βp assumes a 12-stranded β-sheet-rich barrel conformation that spans the bacterial outer membrane (OM), whereby the unfolded passenger domain is translocated from the periplasm to the extracellular milieu through the βp barrel pore (1). OmpB also maintains the βp association within the integral LPS, in the crystalline s-layer surrounding the Rickettsia. A high rate of protein transport (replacement) from the outer membrane to the s-layer maintains the integrity of this protective layer.

## Materials and Methods

### Rickettsia

The EIA panels included 18 SFG species arranged left-to-right according to OmpB sequence (1), plus *Rickettsia felis* (transition species) and SFG LPS. These species were propagated within the BSL3 laboratory in either Vero (CCL-81) or D.mel-2 (CRL-1963), both originally from ATCC. Culture harvests were pooled in liter bottles stored at - 80^0^C until needed for antigen. The frozen material was then thawed within a biological safety cabinet overnight and centrifuged at 2,000 rpm (913 x g) for 7 minutes to pellet cellular debris. Supernates were then pooled and centrifuged at 13,000 rpm (20,440 x g) for 10 minutes. Cell pellets were also saved at -80 °C for later use. Pellets from the high-speed centrifugation were then pooled, resuspended in 10 mL PBS and layered over 30% Renografin (2) for centrifugation at 13,000 rpm. The resulting pellet(s) were washed twice more with PBS, then resuspended in distilled water to elute the s-layer.

This suspension was immediately centrifuged at 13,000 rpm, then this elution procedure was repeated twice. The resulting pooled solution was then centrifuged at 40,000 rpm (20,400 x g) for 120 minutes (50Ti rotor in Beckman L350) to pellet the LPS fraction.

The pellet can be saved as a source of LPS and the soluble protein antigens were treated overnight with Affi-Prep Polymyxin Resin (Bio-Rad) to remove trace amounts of LPS remaining.

In Figures 1-2 these species are abbreviated, and the following (Table 1) is their order on the combined ELISA plates:

**Table 1:** Rickettsia species arranged in columns combining two 96-well ELISA modules.

|  |  |  |
| --- | --- | --- |
| Col 1: *R. africae* | Col 8: *R. japonica* | Col 15: *R. monacensis* |
| Col 2: *R. parkeri* | Col 9: *R. heilongjiangensis* | Col 16: *R. akari* |
| Col 3: *R. siberica* | Col 10: *R. montanensis* | Col 17: *R. australis* |
| Col 4: *R. conorii* | Col 11: *R. raoulti* | Col 18: *R. bellii* |
| Col 5: *R. slovaca* | Col 12: *R. amblyommii* | Col 19: *R. felis* |
| Col 6: *R. honei* | Col 13: *R. rhipacephali* | Col 20: *R. felis* |
| Col 7: *R. rickettsii* | Col 14: *R. helvetica* | Col 21: SFG LPS |

**Figure 1.**
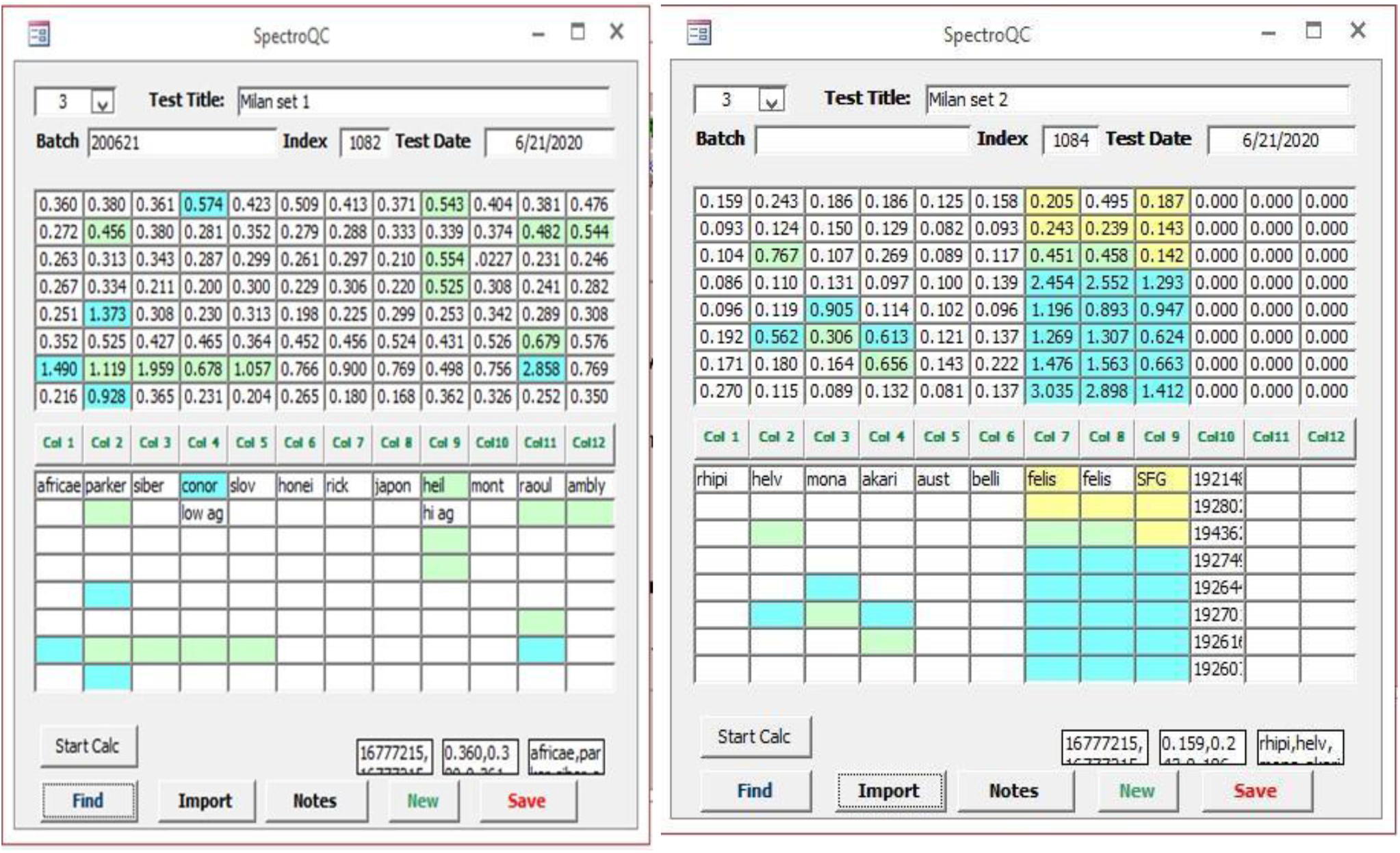
The IgG assay utilized a 1:100 dilution of goat anti-human IgG (Fc)-HRP in Conjugate Diluent (HRP-StabilPlus, Jackson Immunoresearch Laboratories, West Grove, PA) was added for 30 minutes. ELISA Protocol was then followed.

**Figure 2.**
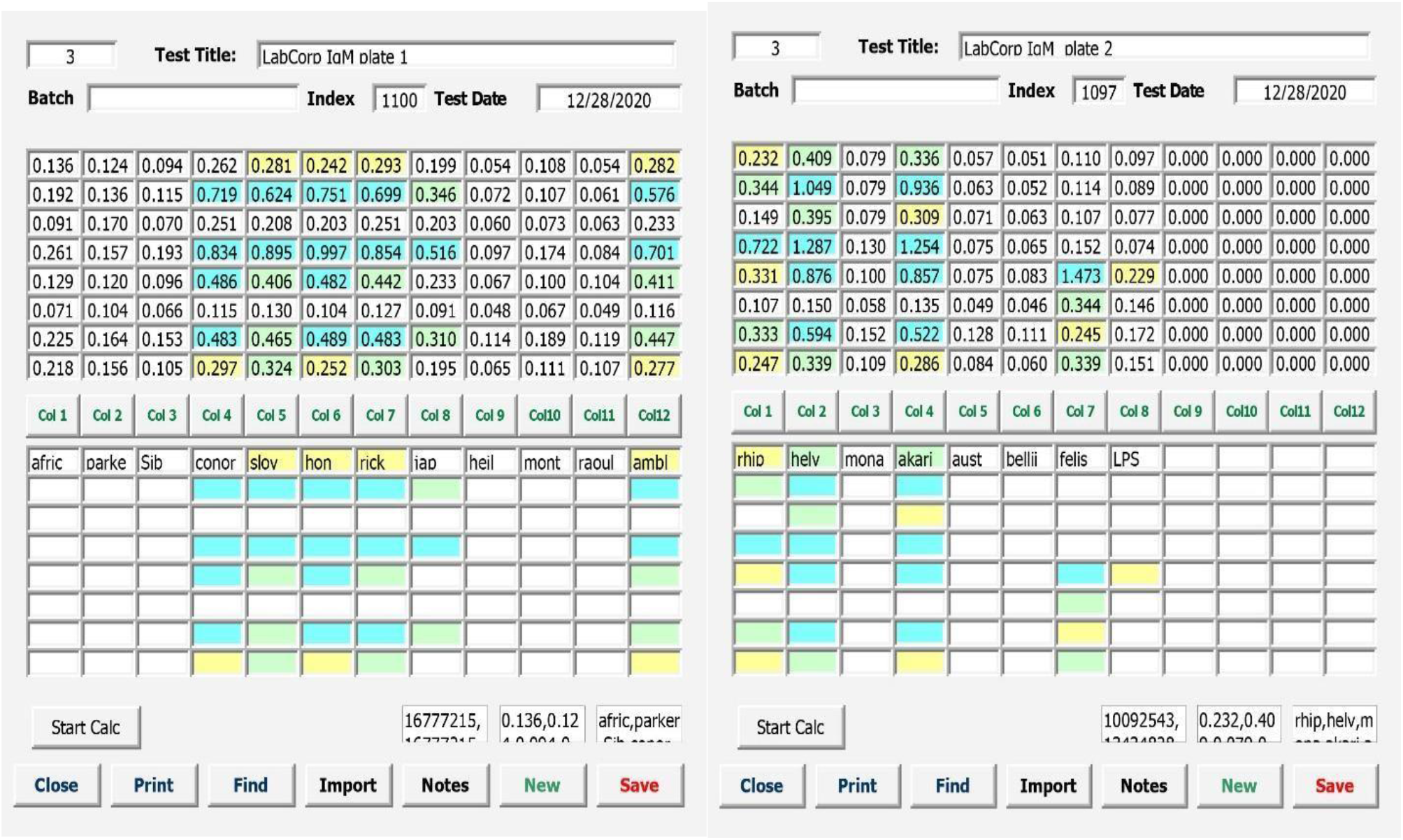
The IgM assay utilized a 1:100 dilution of goat anti-human IgM (Fc)-HRP in HRP-StabilPlus, Jackson Immunoresearch Laboratories) for 30 minutes. ELISA Protocol was then followed.

### Protein Standardization Assay

The three-point calibration curve was performed on antigens according to the manufacturer’s instructions using the included Qubit™ Standard 1–3 reagents. To perform the analysis, 10 µL from each protein sample was combined with 190 µL Qubit™ Working Solution and measured on Qubit Fluorometer (Thermo Fisher Scientific, Waltham, MA, USA). The standard curve was prepared in duplicate and the samples in triplicate. In the end, each protein sample was standardized to 1ug/ml and stored in the walk-in freezer (-10°C to -15°C).

### EIA format

These protein antigens are an artifact of the s-layer elution, each consisting of a twisted heteroduplex of OmpA with OmpB. For use as ELISA antigens they were each diluted to 5 ng/mL in PBS for coating individual 8-well strips. Coating of ELIA modules (Costar Stripwell Microplates) was for 60 minutes at ambient temperature, then directly back-coated with WellChampion (Kementec Solutions A/S, Taastrup, Denmark) for 10 minutes, shaken briefly to remove liquid, then further dried overnight in a low-humidity Drying Room.

### Test sera

Clinical sera were shared by two of our clients without any clinical history or further information. These sera were treated as “unknowns” for different testing series. For IgG testing the sera were from persons living in Europe, while for IgM testing the sera were received as domestic acute-phase. These sera were numbered 1-8 in each group and maintained under refrigeration. They were diluted 1:100 in PBS containing 1% Tween-20 (PBST) for testing.

### ELISA Protocol

Serum rows (100 uL) 1-8 were incubated for 60 minutes at ambient temperature, then washed 3X with PBST. The appropriate HRP-Conjugates were added (100 uL) for 30 minutes at ambient temperature. After a further wash step, TMB (TMB Plus2, Kementec) was added for 10 minutes before Stop Solution (10% NaOH) was added and the plate was read at 450 nm.

## Results

### IgG ELISA Assay

For diagnostic purposes we recommend using purified SFG LPS as antigen, as does the CDC (2). However, we set our panel of 18 antigens to detect the specificity of the individual heteroduplex antigens (see Figure 1).

This IgG reactivity demonstrated species-specificity, especially in convalescent or infections maintained only in immunologic memory. Although this panel is available to customers for differentiation of regional species, this knowledge is basically of academic or epidemiological interest. In acute phase sera these IgG panels also demonstrate a broad peak encompassing near-relation species, while more historic infections produce sharp single-species peaks.

### IgM ELISA Assay

The IgM results were the opposite of the IgG specificity. We suspected that this IgM response was due to a common epitope that had been blocked in almost 50% of the antigen preps by the heteroduplex formation of OmpA with OmpB. These protein chains were formed from linear polypeptides twisted together without native secondary structure while in distilled water. However, these antigens did provide some species-specific IgG reactions when tested in the IgG assay.

### Protein digestion

In our procedure 20 ug from each Rickettsia protein was denatured, reduced and Trypsin digested following the instructions of the manufacturer (Trypsin Platinum VA9000 by Promega Corporation). Digestion of the proteins with Trypsin Platinum (0.5 ug/uL) was carried out overnight at 37^0^C and then terminated with 20% TFA (trifluoroacetic acid).

The resulting trypsin-treated antigens were then assayed on western immunoblot using our reference IgM-positive serum (acute-phase) and antihuman IgM-HRP conjugate. The initial western blot (Figure 3) compares this trypsin treatment with native untreated antigens to locate the IgM-reactive epitope.

**Figure 3.**
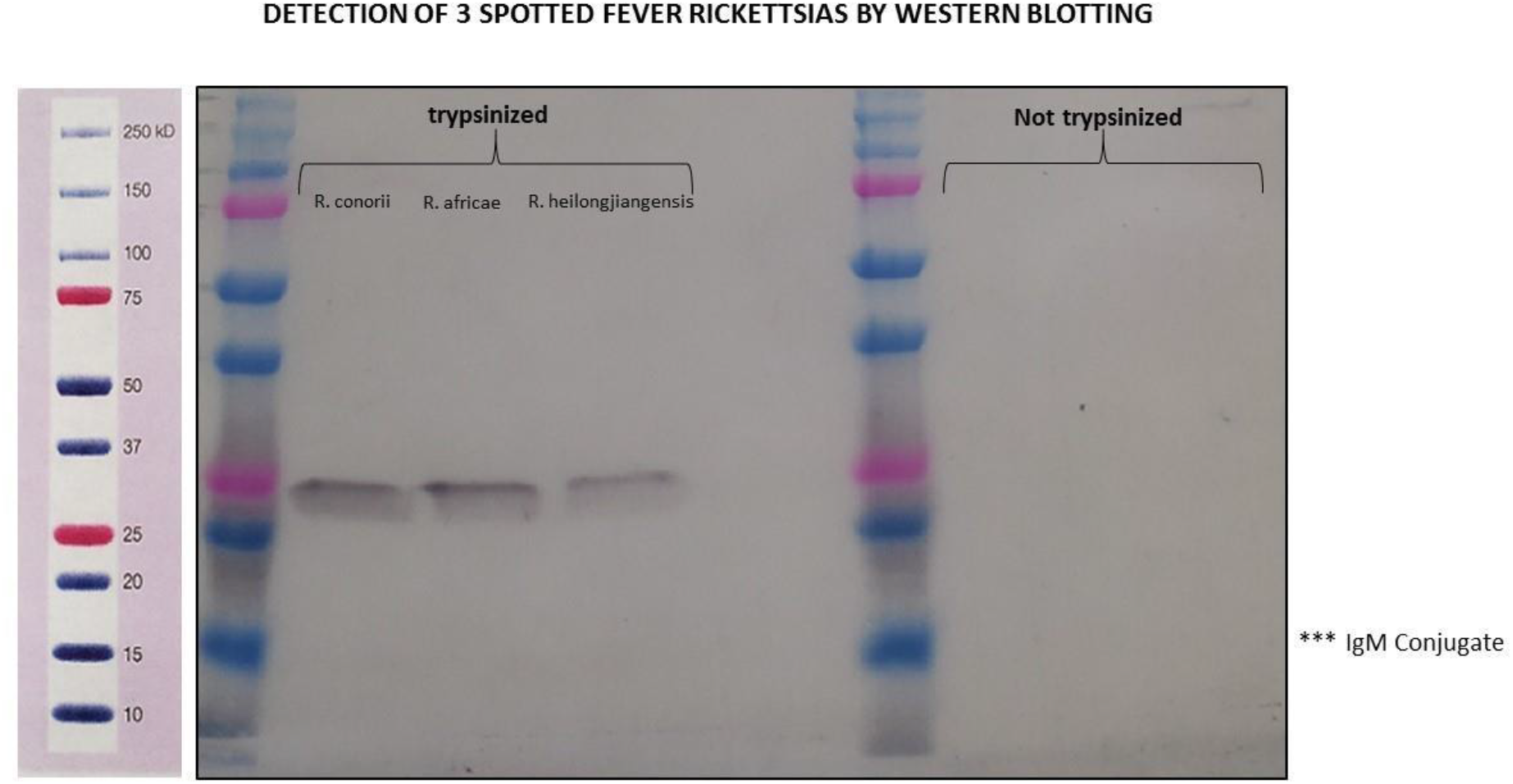
Trypsin treatment shows this reactive fragment to be approximately 25-30 kDa on the trypsin-treated side. The native β-peptide, immediately following enzymatic cleavage from the barrel peptide is 32 kDa (4). The HRP-conjugate here is goat anti-human IgM-specific.

The next western blot (Figure 4) shows seven SFG species, only one (*R. slovaca*) of which was reactive on the original IgM panel of untreated heteroduplexes. This profile suggests that the 25 kDa epitope is related to the β-peptide. When this epitope is concentrated from the trypsin-treated heteroduplexes using a 60 kDa cutoff filter, the EIA reactivity is accurate for detecting immune IgM anti-SFG antibody.

**Figure 4.**
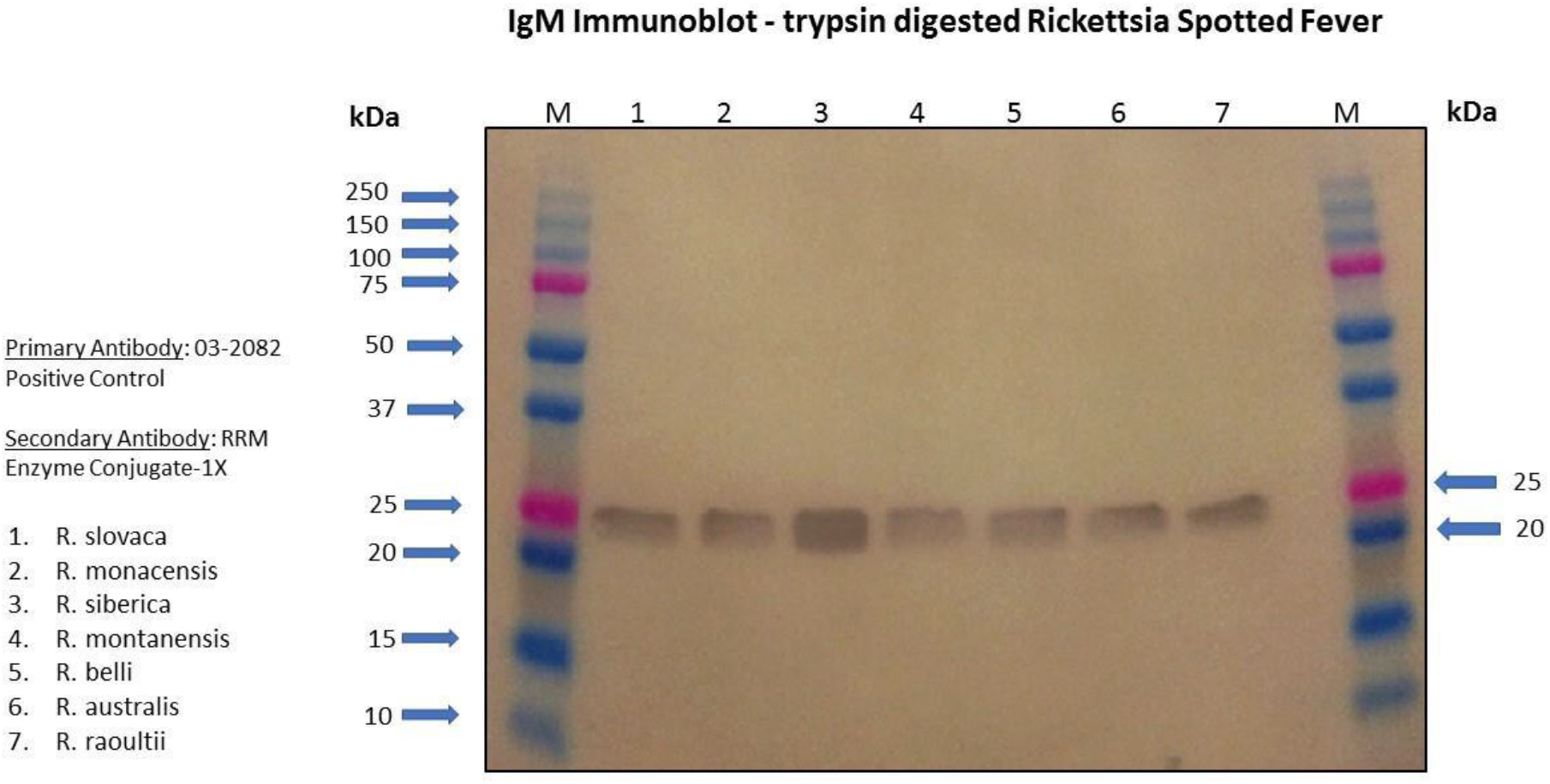
Trypsin treatment shows this reactive fragment to be approximately 25 kDa on all species tested.

## Discussion

We suspect that the native attachment of β-peptide to the OmpB passenger domain allows this epitope to be accessible on both the outer membrane and the outer portion of the s-layer. This may promote survival of the Rickettsia by blocking access to host immune IgG and inhibiting lytic complement attack (5). To maintain this benefit, this epitope is highly conserved across both the SFG species and, to a lesser degree (75%), the TG species.

Earlier (5), an inhibitor of host complement damage, host Complement Factor H was found to bind β-peptide and protect the Rickettsia from complement-mediated damage (death). The further binding of immune IgM puts a second and much larger protein into this small space attached to the β-peptide, although the IgM binding appears to be bound by single binding sites of the pentameric IgM when observed microscopically using anti-IgM Fc specific antisera.

When performing the antigen titration for use in ELISA testing, this β-peptide at 6 increasing dilutions (each dilution 1:10^-8^) was non-reactive with high-titer anti-SFG IgG positive sera using an IgG-specific conjugate. This fact further emphasizes the β-peptide protection against the effects of host immune IgG.

Another recent paper from the Rocky Mountain Laboratory (4) documented a RapL peptidase that cleaves the full 168 kDa protein, freeing the barrel peptide while leaving the β-peptide still embedded in the outer membrane as membrane pore structure. This β-peptide, however, is reactive in our laboratory only after further cleavage from the 32 kDa size to 25 kDa. Our use of trypsin cleaves approximately 81 amino acids from the new N-terminus to activate the reactivity of β-peptide. This cleavage is also required by the 32 kDa recombinant β-peptide. Hence, the unknown peptidase that cleaves these amino acids seems to provide the β-peptide the means to reorient and form a different protein configuration that immediately binds non-covalently to the OmpB barrel peptide.

We plan experimentation and collaboration to determine the reasons this epitope is so important in the evolution and survival of Rickettsia.

## Acknowledgments

We thank Cesar Ordanza, Faezeh Nikravi, Liliane Duraes and Caitlyn Bato for their excellent technical assistance.

